# A possible way to relate the effects of SARS-CoV-2 induced changes in transferrin to severe COVID-19 associated diseases

**DOI:** 10.1101/2022.02.15.480603

**Authors:** Elek Telek, Zoltán Ujfalusi, Gábor Kemenesi, Brigitta Zana, Ferenc Jakab, András Lukács, Gábor Hild

**Author notes:** Correspondence: Gábor Hild M.D., Ph.D.; Department of Biophysics, Medical School, University of Pécs, H-7624, Pécs, Szigeti str. 12, Hungary / / +36 72 / 536-000. These authors contributed to this work equally (Elek Telek, Zoltán Ujfalusi).

## Abstract

The potentially life-threatening Severe Acute Respiratory Syndrome Coronavirus 2 (SARS-CoV-2) infection is responsible for the coronavirus pandemic in 2019 (COVID-19). The transferrin as an essential component of iron-metabolism was suggested to be a link between iron transport associated diseases and COVID-19 infection.

The effect of SARS-CoV-2 on human whole blood was studied by differential scanning calorimetry. The analysis and deconvolution of the thermal transition curves showed that the T_m_ of transferrin related second peak decreased by 5.16 °C (6.4%) in the presence of SARS-CoV-2 virus. The ratio of the under-curve area of the two main peaks was greatly affected while the total enthalpy of the heat denaturation was nearly unchanged in the presence of the virus.

Based on the results it is possible to conclude that SARS-CoV-2 through binding to transferrin can influence it’s Fe^3+^ uptake by inducing thermodynamic changes. Transferrin may stay in iron-free apo-conformational state, which probably depends on the SARS-CoV-2 concentration.

SARS-CoV-2 might induce disturbance in the erythropoiesis due to the free iron overload generated iron toxicity. As a late consequence iron toxicity related hepatocellular carcinoma can even develop.

Our work can support the basic role of transferrin in COVID-19 related severe diseases.

## Introduction

Severe Acute Respiratory Syndrome Coronavirus 2 (SARS-CoV-2) as a member of the Coronavirus family (*Coronaviridae*) is a new human pathogen which is responsible for the coronavirus pandemic started in 2019 (COVID-19) (1). In most cases the SARS-CoV-2 infections cause mild upper respiratory tract symptoms or no symptoms at all. In some patients although life-threatening multi-organ diseases can be observed (1–5). Severe COVID-19 disease has been related to deficient iron metabolism, iron deficiency anaemia (IDA), hypercoagulopathy, thrombosis and ischemic stroke as well (3, 4, 6–9). As a subsequent complication increased incidence of iron toxicity related liver cancer (hepatocellular carcinoma) was also observed (10–13). The relationship between COVID-19 infection and the developed pathological conditions are remained to be elucidated.

Ferritin and transferrin are essential components of the iron metabolism. Ferritin plays a role in iron storage, while transferrin is crucial in iron homeostasis by mediating iron transport to bone marrow where the haemoglobin production takes place which is a key component of the erythropoiesis (14, 15). Increased level of ferritin and transferrin were found in COVID-19 patients and SARS-CoV-2 infected cells as well (16–18). Transferrin may be involved in COVID-19-related IDA, hypercoagulopathy and ischemic stroke as well but the link between transferrin and COVID-19 related dysfunctions is unclear (7, 8, 19, 20). Transferrin is an iron carrier glycoprotein that binds to cellular transferrin receptors and deliver iron by receptor-mediated endocytosis (21, 22).

Transferrin has two lobes which are divided into two structurally different subdomains. Each lobe is able to bind predominantly one iron ion (Fe^3+^) but with different affinity (23). The binding of iron induces conformational change in transferrin. In iron-free state, transferrin has an open-conformation (apo-transferrin), while in its iron-bound form the conformation is more closed (holo-transferrin) (24). Transferrin can be actively involved in the coagulation cascade as a procoagulant by inhibiting antithrombin and factor XIIa (25). A recent study showed that the expression of transferrin increased by age and its level was consequently higher in males. As the antithrombin level was not affected by age or gender the ratio of transferrin/antithrombin is the highest in elder males (20). These data are in agreement with the increased risk of severe COVID-19 disease found in elderly males (26). Based on these considerations increased transferrin expression and transferrin/antithrombin ratio might have significant contribution to COVID-19-related coagulopathy with more severe outcome in older male patients. To study the effect of SARS-CoV-2 on human whole blood calorimetric measurements were performed to elucidate the molecular adjustment of transferrin to the viral invasion.

## Material and methods

### Sample preparation

Blood was taken in light blue topped BD Vacutainer® tubes treated with anticoagulants by healthcare professionals from healthy volunteers and kept on ice. The applied procedures were approved by the Local Ethical Committee of the UP MS (Certificate No. 8549-PTE2020). SARS-CoV-2 virus isolation was performed in a Biosafety level 4 (BSL-4) laboratory of the Szentágothai Research Centre, University of Pécs, Hungary. During the experimental procedure we used a Hungarian SARS-CoV-2 isolate (isolated on Vero E6 cells, GISAID accession ID: EPI_ISL_483637). Infection of human whole blood was also carried out in the BSL-4 laboratory by mixing 50 µl of isolated SARS-CoV-2 virus in a concentration of 5.62×10^6 TCID50 with 850 µl of whole blood sample. Thereafter, 800 µl of the mixture was pipetted directly into a conventional hastelloy batch vessel (V_max_ = ∼1 mL) and the samples were incubated at either room temperature (24 °C), or at body temperature (37 °C) or at 40 °C for 2, 15, 25 or 50 hours. The properly sealed batch vessels (containing the samples) have been cleaned by sterilizing agent (mikrozid® AF liquid - Schülke & Mayr GmbH) and kept in it until the DSC analysis could be performed.

### DSC measurements

Differential scanning calorimetry (DSC) is a widespread thermoanalytical technique to study the thermodynamic properties of clinical samples (27–31). To investigate the effect of SARS-CoV-2 on the human whole blood thermal denaturation measurements were performed with SETARAM Micro DSC-III calorimeter. Each measurement was performed in the range of 20 – 100 °C by using 0.3 K·min^-1^ heating rate. The sample and the reference (normal saline) were balanced with precision of ± 0.05 mg in order to avoid corrections with the heat capacity of the vessels. A second thermal scan of the denatured sample was carried out for baseline correction. Deconvolution of DSC data, the analysis of the melting temperature (T_m_) and the calculation of the relative enthalpy change from the area under the deconvolved heat absorption curves were analysed by using OriginLab Origin®2021 software.

## Results & Discussion

To demonstrate the effect of SARS-CoV-2 virus on blood components, calorimetric measurements were performed on anticoagulated human whole blood samples in the presence and absence of SARS-CoV-2. The virus caused significant change in the thermodynamic properties of blood components. As human blood contains relatively high number of different proteins the resulted endothermic denaturation peaks are the collective representations of the individual components (Fig. 1 and 3). To overcome this issue, we deconvolved the main two peaks and treated them separately (Fig. 2 and 4). The first large endothermal peak (Peak 1) is mainly due to the presence of albumin along with some other proteins as well (Fig. 1-4), while the second large peak (Peak 2) is typically the combination of γ-globulin (IgG) and the β-globulin type transferrin (Fig. 1-4) (32) alongside with some extra proteins which are present at relatively low concentration.

**Figure 1.**
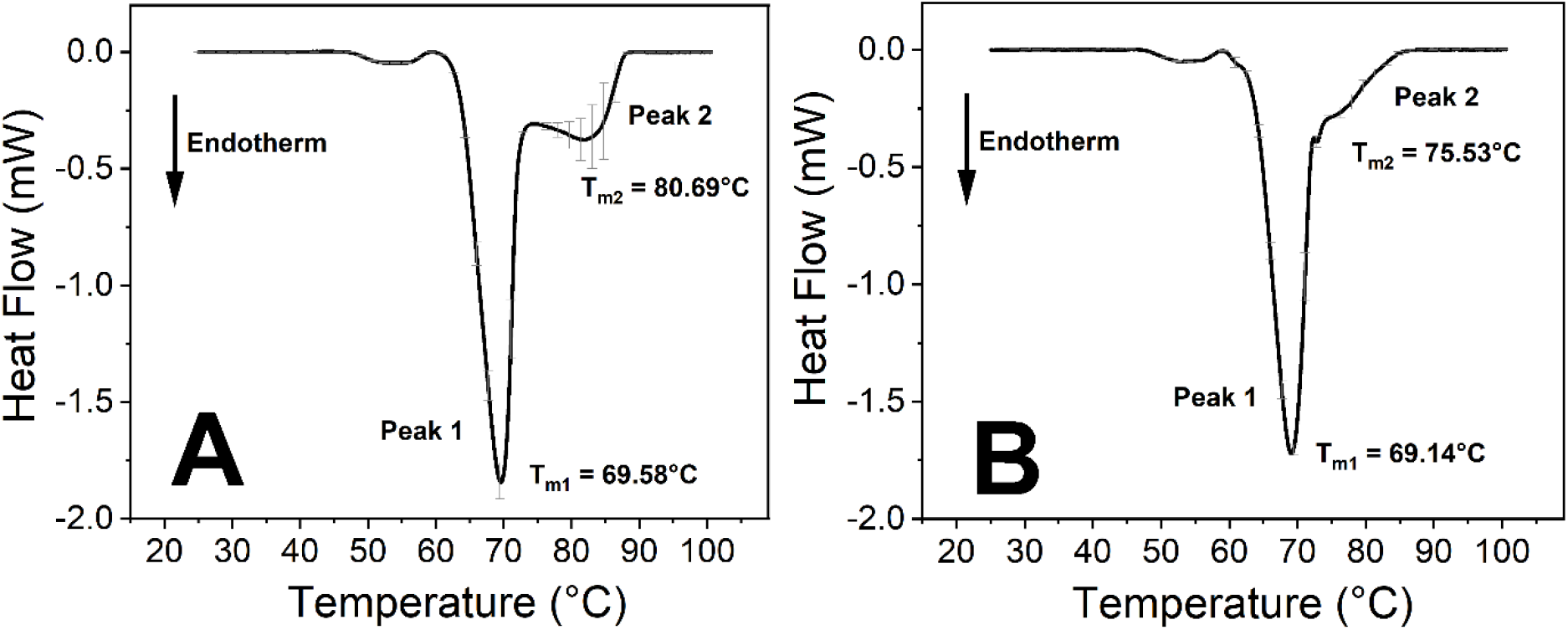
Thermal analysis of untreated and SARS-CoV-2 - treated anticoagulated human whole blood samples. **(A)** Control 25 hours incubation at 37 °C. **(B)** Treated with SARS-CoV-2 and incubated for 25 hours at 37 °C. The measurements were performed at least four times independently (n ≥ 4) and the results are presented as mean ± SD.

**Figure 2.**
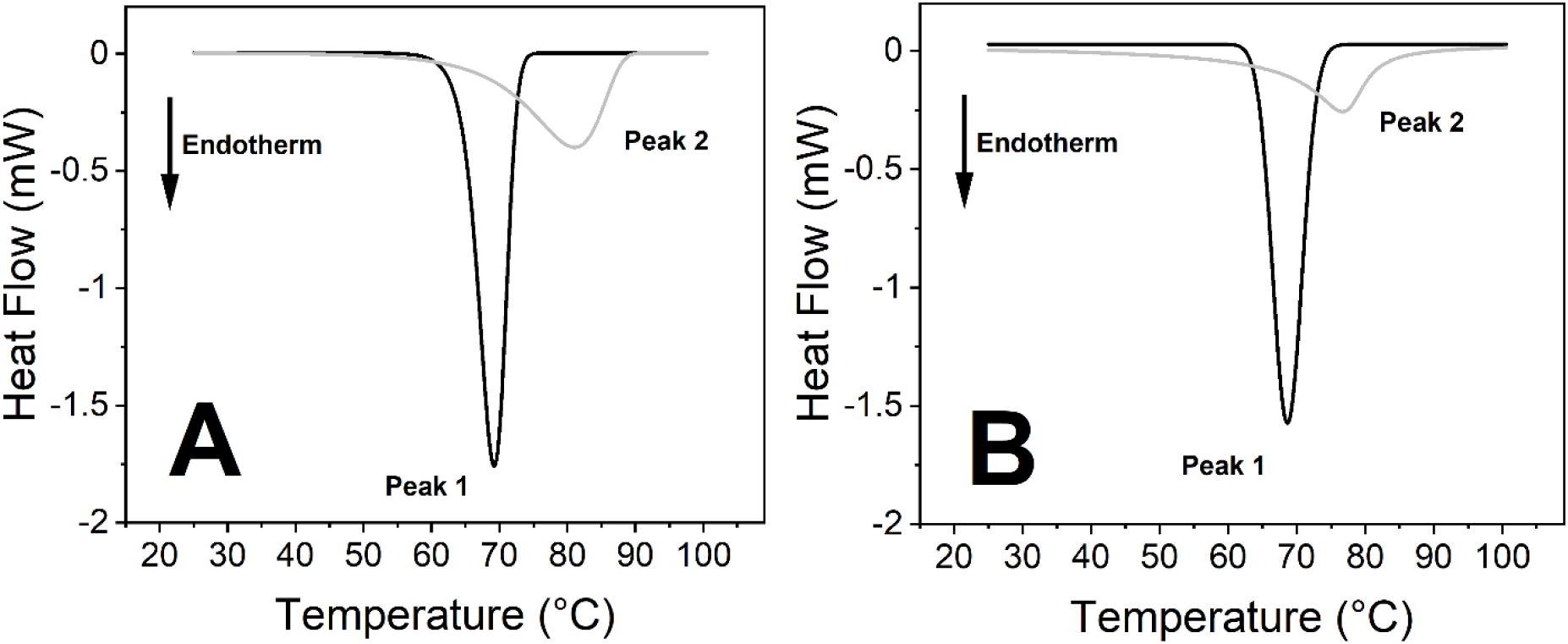
Deconvolution of Peak 1 and Peak 2 from Fig. 1 DSC plots. **(A)** Control 25 hours incubation at 37 °C. **(B)** SARS-CoV-2 treatment incubated for 25 h at 37 °C.

**Figure 3.**
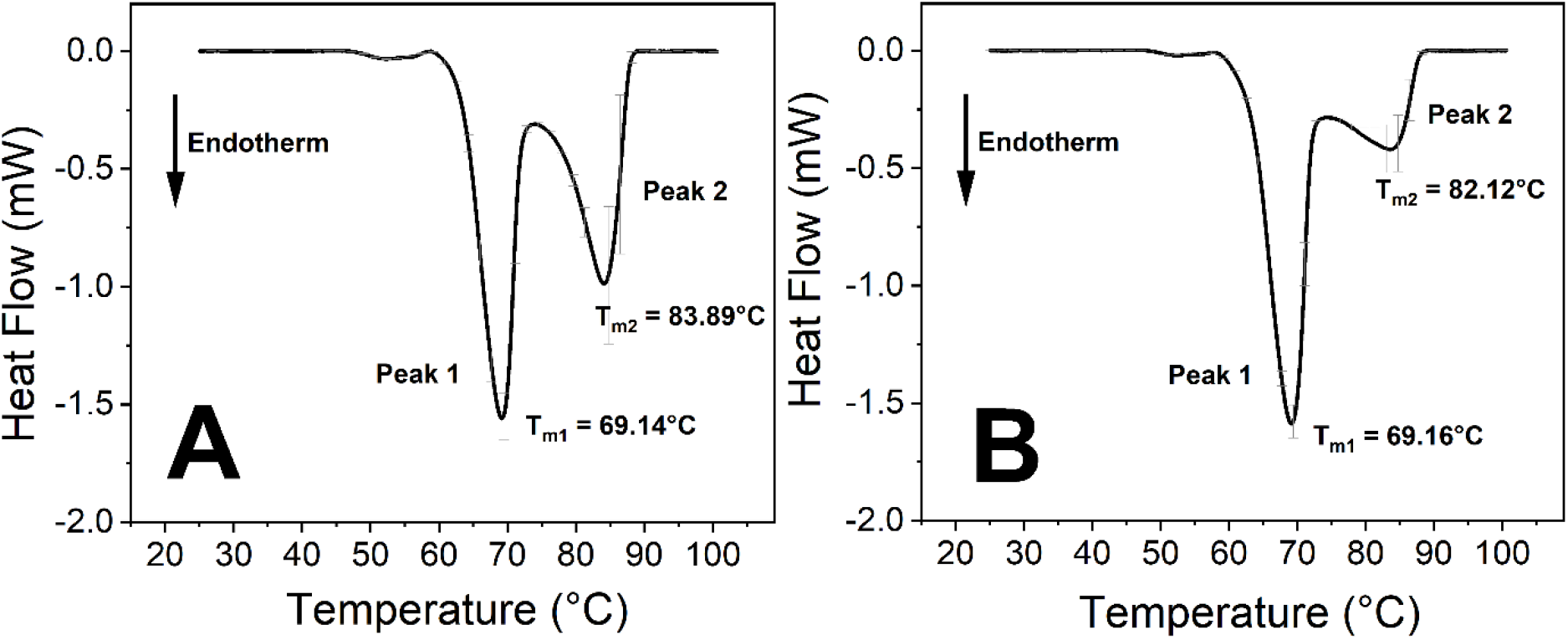
Thermal analysis of untreated and SARS-CoV-2 - treated human anticoagulated whole blood samples. **(A)** Control 50 hours incubation at 37 °C. **(B)** Treated with SARS-CoV-2 and incubated for 50 hours at 37 °C. The DSC data represent the mean ± SD of at least four independent measurements (n ≥ 4).

**Figure 4.**
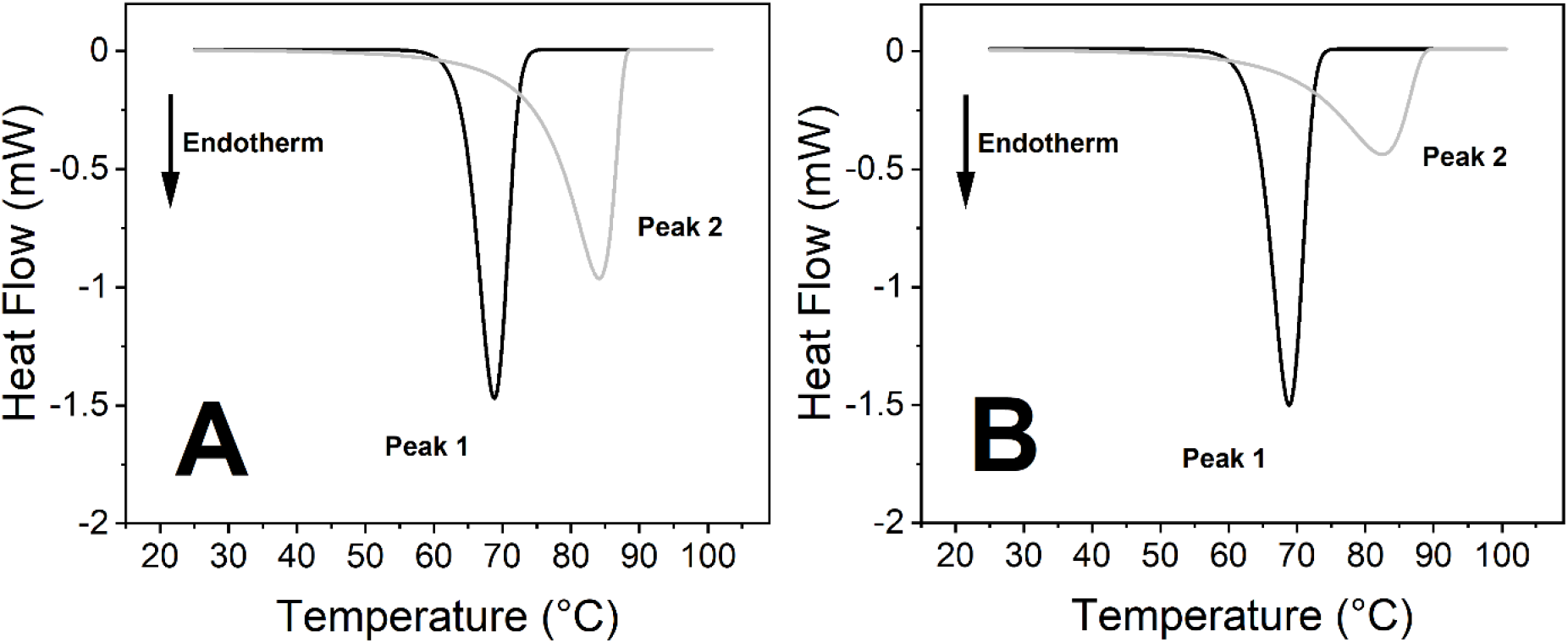
Deconvolution of Peak 1 and Peak 2 from Fig. 3 DSC plots. **(A)** Control 50 hours incubation at 37 °C. **(B)** Treated with SARS-CoV-2 and incubated for 50 hours at 37 °C.

The incubation of the samples was performed at different temperatures to see the temperature sensitivity of the virus activity at room temperature (24 °C), body temperature (37 °C) and at a high fever simulating temperature (40 °C) as well. After a 2 hours long incubation time at different temperatures the shape and the main thermal parameters of the control curves (Fig. 6) and the treated samples were the same (Fig. S1). Long incubations times (50 hours) at all three temperatures show indisputable effects of the SARS-CoV-2 virus on the blood components (Fig. 3 and Fig. S2) while on a shorter time scale (25 hours) only the 37 °C incubation temperature proved to be efficient. According to these results we focused on the effects of SARS-CoV-2 caused on human blood samples at 37 °C. At incubation times less than 25 hours there have been no significant difference between the control and the virus-containing samples, therefore 25 hours and 50 hours incubation time intervals were selected to explore the thermal properties of the samples (Fig. 1-5).

**Figure 5.**
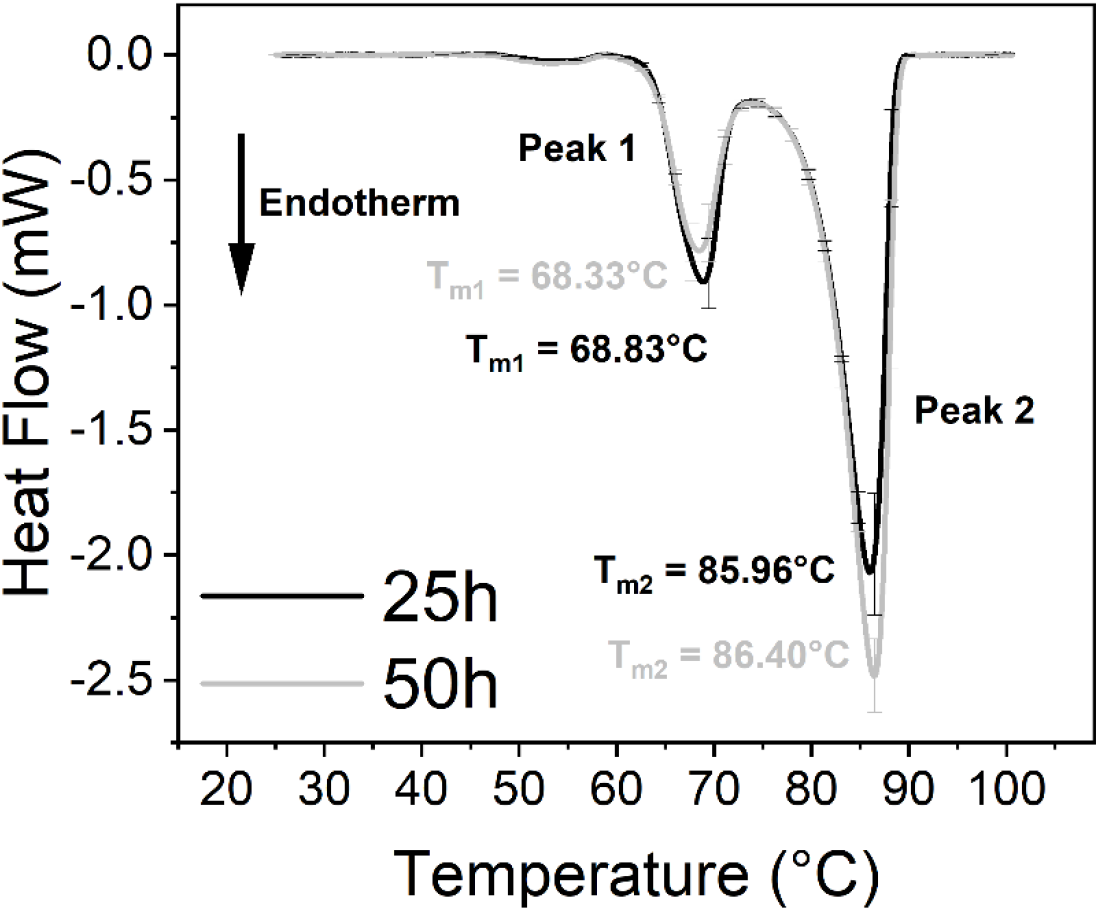
Anticoagulated human whole blood treated with DMEM (Dulbecco’s Modified Eagle Medium) as a control at 37 °C. The measurements were performed three times independently (n = 3) and the results are presented as mean ± SD.

**Figure 6.**
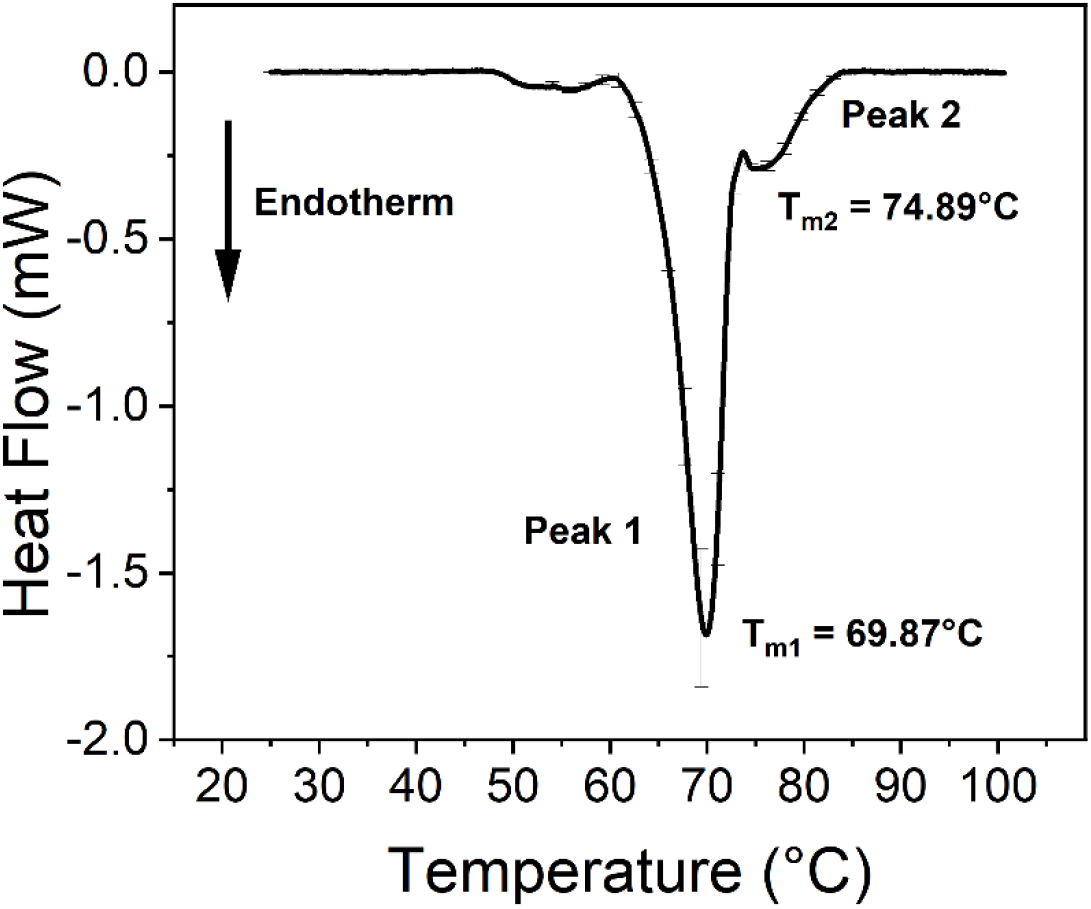
Thermal analysis of untreated anticoagulated whole blood samples after 2 hours of incubation. The plot is an average of the same measurements repeated at different incubation temperatures (24 °C, 37 °C and 40 °C). The DSC data represent the mean ± SD of three independent measurements (n = 3).

There is no significant difference in T_m_ values (melting temperature – peak of the endothermic curve where ∼50% of the proteins are denatured) of the first peaks (Peak 1) at any incubation time between the treated and control samples (Fig. 1-5 and Tables 1-2). Interestingly other characteristic parameters of Peak 1 like overall shape, half-width and area remained unchanged as well. Unlike Peak 1, Peak 2 is very sensitive for both the incubation time and the presence of the SARS-CoV-2 virus as well. The ratios of the integrated area of Peak 2 to Peak 1 change dramatically with the increasing incubation time (Fig. 1-4 and Table 1). Mainly the undercurve area and T_m_ of Peak 2 fluctuates while Peak 1 remains constant (Fig. 1-4). These alterations of the second peak could be emphasized by the deconvolution of the main peaks (Fig. 2 and 4) and thus the difference between the virus-treated and control samples is conspicuous. The huge differences between the virus-treated and control samples at longer incubation times suggest that the undisturbed consistency of the treated samples are effects of SARS-CoV-2 infection where Peak 2 plays a key role.

**Table 1.** Detailed thermal characterisation of the primary and deconvolved DSC data of human whole blood in the absence and presence of SARS-CoV-2. (* determined from the deconvolution of averaged data). Data represent mean ± SD (n ≥ 4).

| Thermal parameters of Peak 1 and 2 from DSC data |  |  |  |  |
| --- | --- | --- | --- | --- |
|  | 25 hours incubation |  | 50 hours incubation |  |
|  | untreated | SARS-CoV-2 | untreated | SARS-CoV-2 |
| $T_m$ of Peak 1 (°C) | $69.58 \pm 0.11$ | $69.14 \pm 0.14$ | $69.14 \pm 0.10$ | $69.16 \pm 0.09$ |
| $T_m$ of Peak 2 (°C) | $80.69 \pm 1.36$ | $75.53 \pm 0.24$ | $83.89 \pm 0.58$ | $82.12 \pm 1.23$ |
| Peak 2:1 ratio * | 0.63 | 0.54 | 1.25 | 0.74 |
| Calculated $\Delta H$ (J/g) | $3.51 \pm 0.13$ | $2.85 \pm 0.05$ | $4.12 \pm 0.19$ | $3.44 \pm 0.10$ |

**Table 2.** Detailed thermal characterisation of the primary and deconvolved DSC data of control measurements in the presence of DMEM. (* determined from the deconvolution of averaged data). Mean ± SD (n = 3).

| Thermal parameters of Peak 1 and 2 from DSC data of DMEM control measurements |  |  |
| --- | --- | --- |
|  | 25 hours incubation | 50 hours incubation |
| $T_m$ of Peak 1 (°C) | $68.83 \pm 0.17$ | $68.33 \pm 0.28$ |
| $T_m$ of Peak 2 (°C) | $85.96 \pm 0.24$ | $86.40 \pm 0.18$ |
| Peak 2:1 ratio * | 3.30 | 4.25 |
| Calculated $\Delta H$ (J/g) | $4.31 \pm 0.06$ | $4.64 \pm 0.04$ |

The main component of Peak 2 is transferrin. Transferrin has a crucial role in the distribution and regulation of iron (mainly in the form of Fe^3+^) throughout the whole body. It binds iron with very high affinity, which gives its main function as to reduce the free iron content of the blood. Moreover, transferrin prevents the reduction of the bound Fe^3+^ and thus Fe^2+^ caused toxicity (23). One transferrin can bind two Fe^3+^ ions in a strict order. The first metal ion is always bound to the C-terminal of the protein while the second one in the cleft between the lobes on the N-terminal part of transferrin (33). The thermal properties of this protein are strongly dependent on its iron-bound state. Binding one metal ion increases the melting temperature significantly and the incorporation of a second Fe^3+^ shifts the T_m_ of transferrin to even higher temperatures (23).

We got the most consistent results at 37°C which is roughly the normal body temperature. Between 2-15 hours of incubation there were not any significant difference between the virus-treated and the control blood samples (Fig. 6, Fig. S1 and Fig. S3). The drastic changes developed after 15 hours of incubation time (Fig. 1 and 3). The melting temperature of Peak 2 increased significantly for the non-treated samples from 80.69 °C (25 hours) to 83.89 °C (50 hours) which suggests that transferrin bound more Fe^3+^ ions which are probably released from some other blood sources. The undercurve area also increased significantly in case of the control samples (Fig. 1A, 2A, 3A, 4A and Fig. 6) indicating that more energy is required to denature the protein. Based on these results after the conformational change the transferrin ended up in a more stable configuration.

The enthalpy (ΔH) of Peak 2 decreased in case of the SARS-CoV-2 virus-treated samples compared to the control (Table 1). The presence of SARS-CoV-2 seems to prevent the development of the stabilisation of transferrin, which clearly suggest that the virus interacts with transferrin and seems to inhibit the uptake of Fe^3+^ by the protein. We observed the same tendency in enthalpy change when the incubation temperature was 24 °C and 40 °C but the development of the structural change of transferrin developed after longer incubation times only (data not shown).

We used VeroE6 (ATCC No. CRL-1586) type cells to replicate SARS-CoV-2 in DMEM (Dulbecco’s Modified Eagle Medium), so a strict test of the effect of this medium was necessary to be able to exclude the effects of DMEM (cell debris was removed by centrifugation – see Materials and Methods). In these experiments the medium was added to the anticoagulated blood samples only and after the given incubation times the experiments were completed (Fig. 5, Table 2). In the presence of DMEM the T_m_ values shifted to a higher level with the increased level of the ΔH values (Table 1 and 2.). The results clearly show that DMEM alone has a very strong and opposite effect in the lack of virus which makes the effect of SARS-CoV-2 on the blood components even more dramatic. Control experiments have also been made with the SARS-CoV-2 virus under the above-described conditions but with saline solution as a reference. These results provided flat, horizontal DSC curves with negligible disturbances (data not shown) which suggest that the presence of SARS-CoV-2 alone does not have any contribution to the DSC signals at all.

## Conclusion

The 2019 pandemic caused by acute respiratory syndrome coronavirus 2 (SARS-CoV-2) can cause mild upper respiratory tract symptoms and life-threatening multi-organ diseases as well (1–5). Severe COVID-19 disease may disturb the iron metabolism and can cause iron deficiency anaemia (IDA), hypercoagulopathy, thrombosis, ischemic stroke or even iron toxicity-related hepatocellular carcinoma (HCC) as a late complication (3, 4, 6, 7, 9–11). Both, ferritin and transferrin have a key role in the iron metabolism. Ferritin is crucial in iron binding and storage (14) while transferrin is important in iron homeostasis by transporting iron for erythropoiesis (15, 16). In COVID-19 patients and SARS-CoV-2 infected cells increased level of ferritin and transferrin was found (17–19) suggesting the involvement of these iron metabolism-related proteins in the development of the disease.

Transferrin might have a role in IDA, hypercoagulopathy and ischemic stroke as well (7, 8, 20). Although there seems to be a relationship between transferrin and COVID-19 infection the evidence based real pathological background is still unknown. In order to study the effect of SARS-CoV-2 on blood components calorimetric measurements were carried out at different temperatures (24 °C, 37 °C, 40 °C) after 25 and 50 hours incubation time with anticoagulated human whole blood samples in the absence and presence of the virus. Initially the samples were prepared at 37 °C incubation temperature to simulate the effect of SARS-CoV-2 at body temperature.

DSC measurements carried out after 25 hours long incubation at 37 °C showed no significant change in the T_m_ of Peak 1 in the absence and presence of SARS-CoV-2 (Fig. 1 and 2, Table 1). The thermal analysis of Peak 2 revealed a substantial 6.4% decrease of the T_m_ (ΔT_m_= 5.16 °C) and 18.80% decrease in the ΔH value when the samples were treated with the virus (Fig. 1B and 2B, Table 1).

When the untreated sample were incubated for 50 hours at 37 °C, the T_m_ of Peak 1 was nearly the same as after 25 hours incubation and it was independent from the presence of the virus. The T_m_ of Peak 2 of the untreated sample shifted to 83.89 °C (3.96% increase) while the under-curve area of Peak 2 increased to 4.12 J/g (17.38% increase) compared to the samples incubated for 25 hours. The T_m_ of Peak 2 of the virus treated samples treated for 50 hours long was nearly the same as the untreated one while the ΔH value decreased by 16.5% in the presence of SARS-CoV-2 which is the same tendency that was observed after shorter incubation time (Fig. 3B and 4B, Table 1).

These changes of the thermal parameters can suggest that a more compact and stable conformation of Fe^3+^-bound transferrin was formed after a longer incubation time in the presence and absence of the virus as well. Over 50 hours incubation at 37 °C probably more Fe^3+^ can be released from haemoglobin and able to bind transferrin, therefore the concentration of Fe^3+^-bound holo-transferrin is increased. As transferrin has two binding sites for Fe^3+^ by the increased incubation time the saturation of both of the binding sites increases that results in a higher under curve area and T_m_ value for the untreated and treated samples as well.

At shorter incubation time the T_m_ and ΔH of Peak 2 decreased as well while at longer incubation time only the ΔH decreased in the presence of the SARS-CoV-2. All these changes clearly show the significant sensitivity of Peak 2 for the presence of SARS-CoV-2 (Table 1). The virus undeniably decreases the thermal stability of transferrin. As a consequence, the conformation becomes less stable, and the energy consumption of the denaturation is lower compared to the untreated blood samples. The thermal denaturation can be initiated more easily and with less energy due to the presence of the SARS-CoV-2 virus which can be really pronounced after shorter incubation time. When the samples were treated too long the T_m_ value became insensitive for the presence of SARS-CoV-2 (Table 1).

Control measurements were performed to exclude the effect of the virus buffer (DMEM) on the human whole blood components. After treating the samples with DMEM exclusively the analysis showed no significant change in the T_m_ of Peak 1 and Peak 2 while the under-curve area was significantly higher after 25 and 50 hours as well compared to the samples where the DMEM was present in smaller amount. The opposite tendency in the change of ΔH in the absence of the virus can suggest an even larger effect of SARS-CoV-2 on transferrin and confirms our experimental findings (Fig. 5, Table 2).

Our calorimetric measurements revealed that SARS-CoV-2 influences the thermodynamic properties of transferrin (Peak 2) suggesting that the virus can interact with transferrin and probably inhibits the uptake of Fe^3+^ (Fig. 7). After shorter incubation time (25 hours) the less rigid iron-free apo-transferrin seems to be predominant with significantly lower melting temperature and lower under curve area as well. After longer incubation time (50 hours) probably the concentration of Fe^3+^-bound holo-transferrin is increased as more Fe^3+^ is available after its release from haemoglobin, therefore the area under the curve area became higher in the control samples. The effect of SARS-CoV-2 can counterbalance at least partially the uptake of Fe^3+^ which can develop only after a longer impact time. The Fe^3+^-bound holo-transferrin can be predominant after long incubation and SARS-CoV-2 has enough time to compensate at least partially for its presence. Consequently, change in the T_m_ cannot be observed since the holo-transferrin level is supposedly more abundant and it can also mask the interaction of SARS-CoV-2 with apo-transferrin. Our findings are in accordance with previously published data, where DSC measurements of iron-titrated transferrin resulted in similar T_m_ values depending on the occupied Fe^3+^ binding sites (23).

**Figure 7.**
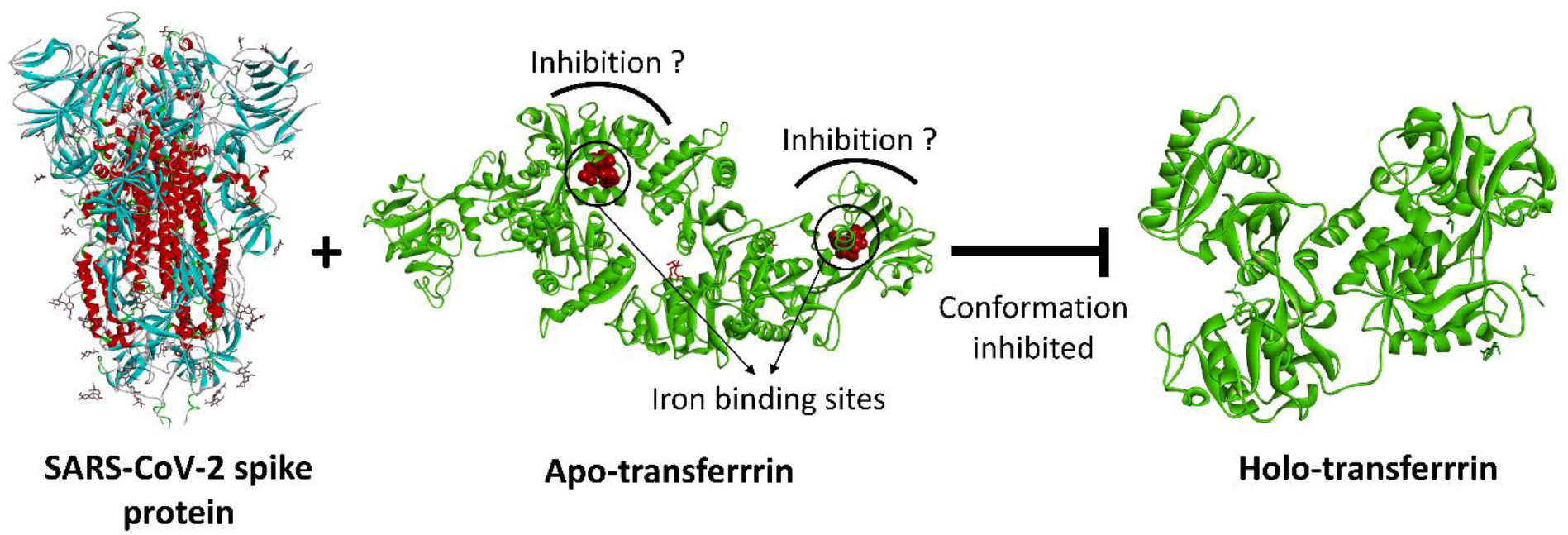
Schematic representation of how SARS-CoV-2 (PDB code: 7jwy) may blocks Fe^3+^ uptake of apo-transferrin (PDB code: 2hau) leading to inhibition of holo-transferrin (PDB code: 3v83) conformation.

As a conclusion, SARS-CoV-2 may influence the transferrin-mediated iron transport processes leading to inefficient erythropoiesis causing IDA followed by iron toxicity due to the free iron overload (34). In addition, HCC can also be developed as a late complication of iron toxicity-induced ROS production caused indirectly by SARS-CoV-2 (10, 11, 13). Moreover, the increased level of transferrin and the change in transferrin/antithrombin ratio (20, 26) may cause misregulation of the coagulation system, especially in males leading to COVID-19-related hypercoagulopathy, thrombosis and ischemic stroke. The schematic model of the effect of SARS-CoV-2 is represented on Fig. 8. It is also known that Covid infection can directly influence the oxygen uptake by destroying the lung tissue and inducing long term fibrosis in the lung parenchyma as well (35, 36).

**Figure 8.**
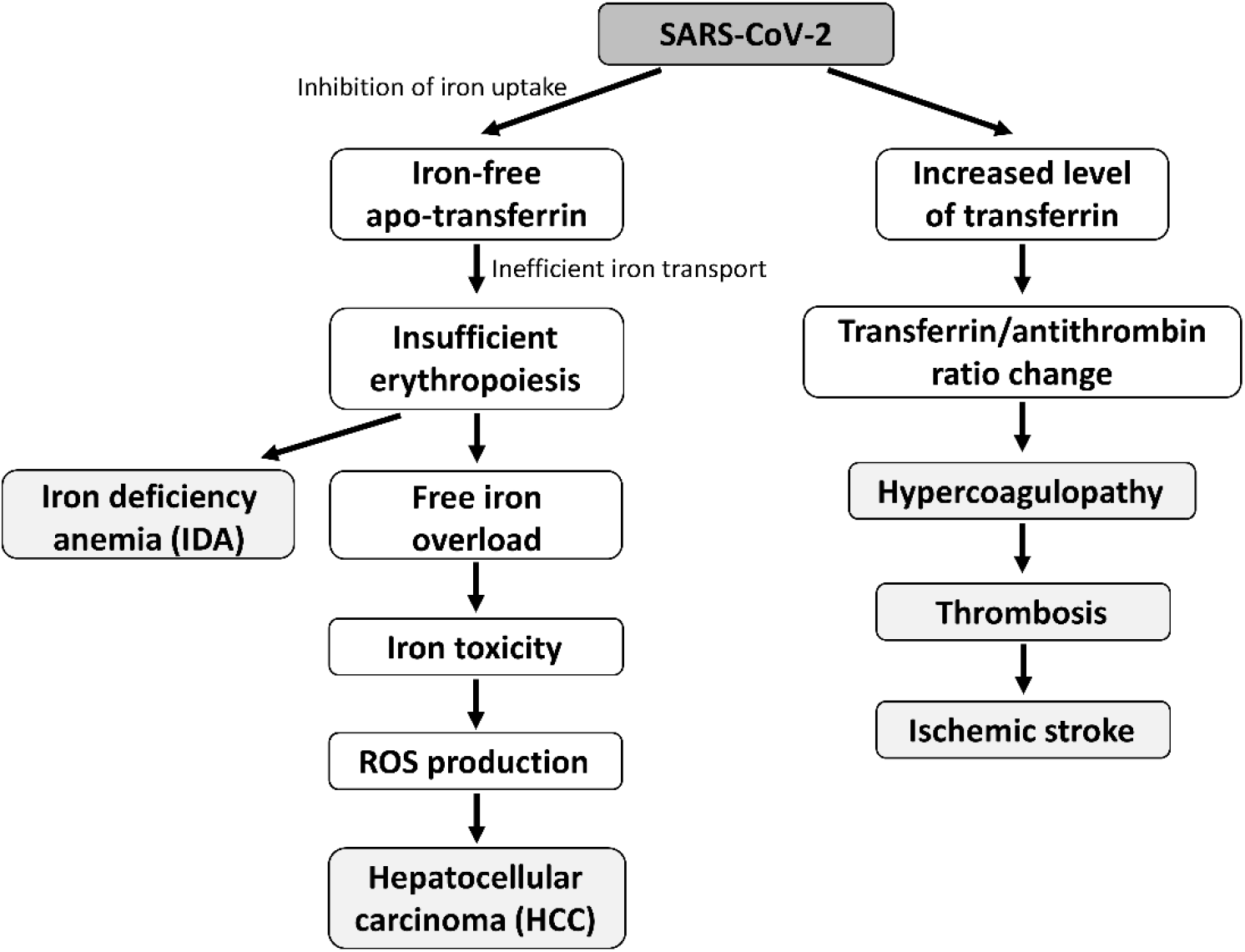
Schematic model of the effect of SARS-CoV-2 on transferrin emphasizing the relationship with COVID-19-related severe diseases. The left flow chart represents new insights of possible SARS-CoV-2 influence on transferrin by inhibiting Fe^3+^ uptake based on our thermal analysis findings may leading to IDA and HCC. The right flow chart of the model summarizes how the increased transferrin level change the transferrin/antithrombin ratio (25) probably causing hypercoagulopathy, thrombosis and ischemic stroke.

Based on these data it seems that SARS-CoV-2 can attack not only on different organs but on different cellular and molecular levels as well which potentially makes it a highly effective and dangerous enemy in the war fought against it.

## Author contributions

Conceptualization: G.H., E.T., Z.U.; Experiment design: G.H., E.T., Z.U.; Experiment performance and analysis: Z.U. & E.T.; Virus isolation, sample preparation: G.K., B.Z.; Writing first draft of the paper: E.T. & Z.U.; All authors contributed to the final version of the manuscript.

## Acknowledgements

This work was supported and funded by UP MS grant EFOP-3.6.3-VEKOP-16-2017-00009 (Z.U.) and UP MS grants KA-2020-09 (Z.U.), KA-2019-40 (belongs to G.H. and Z.U.).

