## Supplementary materials for "A possible way to relate the effects of SARS-CoV-2 induced changes in transferrin to severe COVID-19 associated diseases"

### Supplementary figures

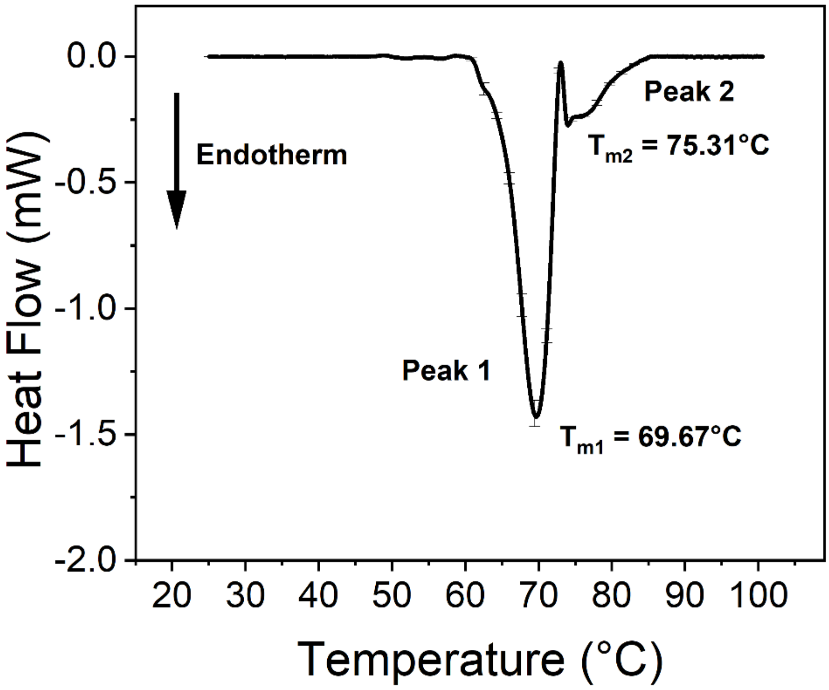


**Figure S1:** Thermal analysis of virus-treated anticoagulated whole blood samples after 2 hours of incubation. The plot is an average of the same measurements repeated at different incubation temperatures (24 °C, 37 °C and 40 °C). The DSC data represent the mean ± SD of three independent measurements (n = 3).

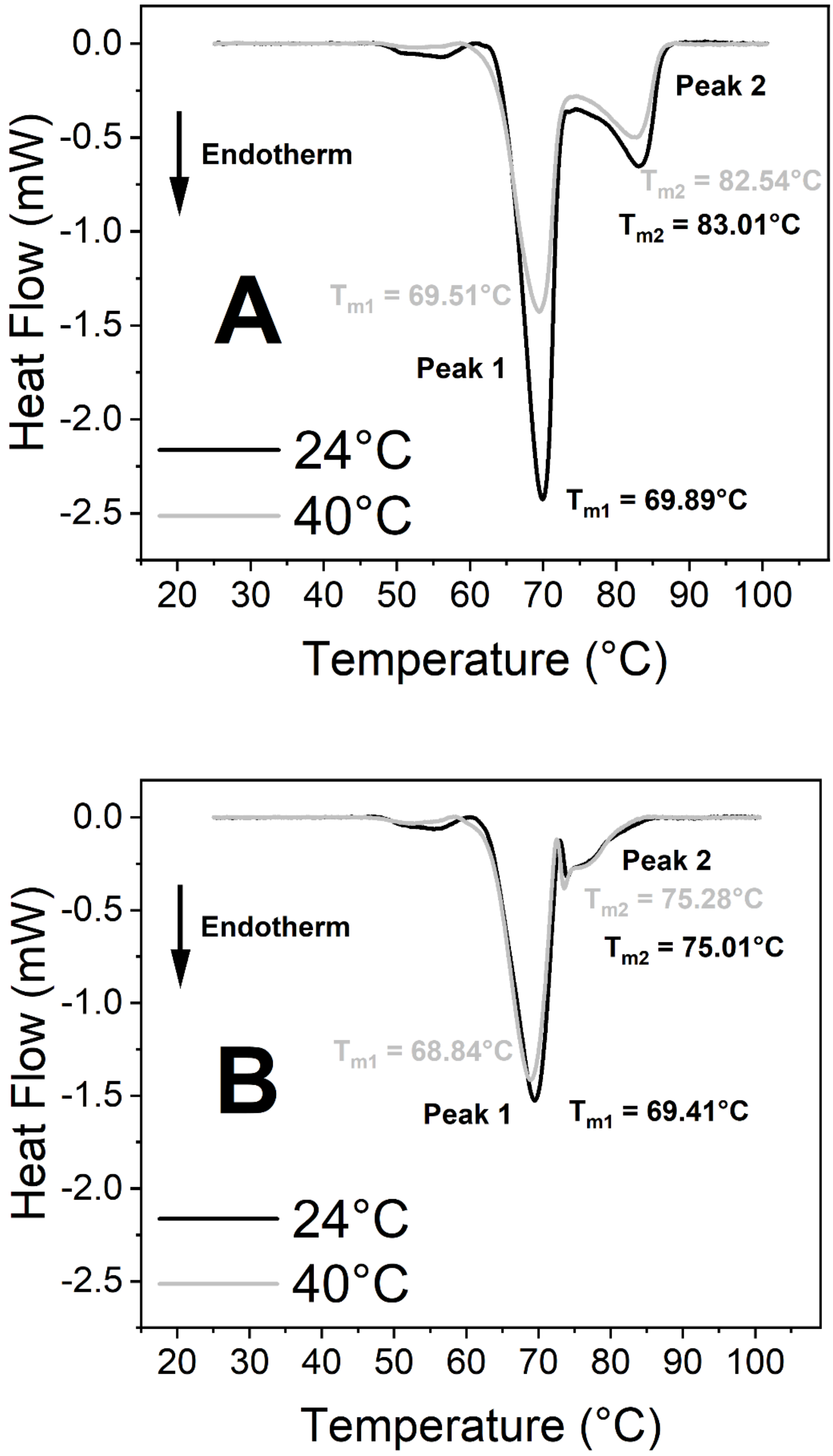


**Figure S2:** Thermal analysis of control (A) and virus-treated (B) anticoagulated whole blood samples after 50 hours of incubation at 24 °C and at 40 °C.

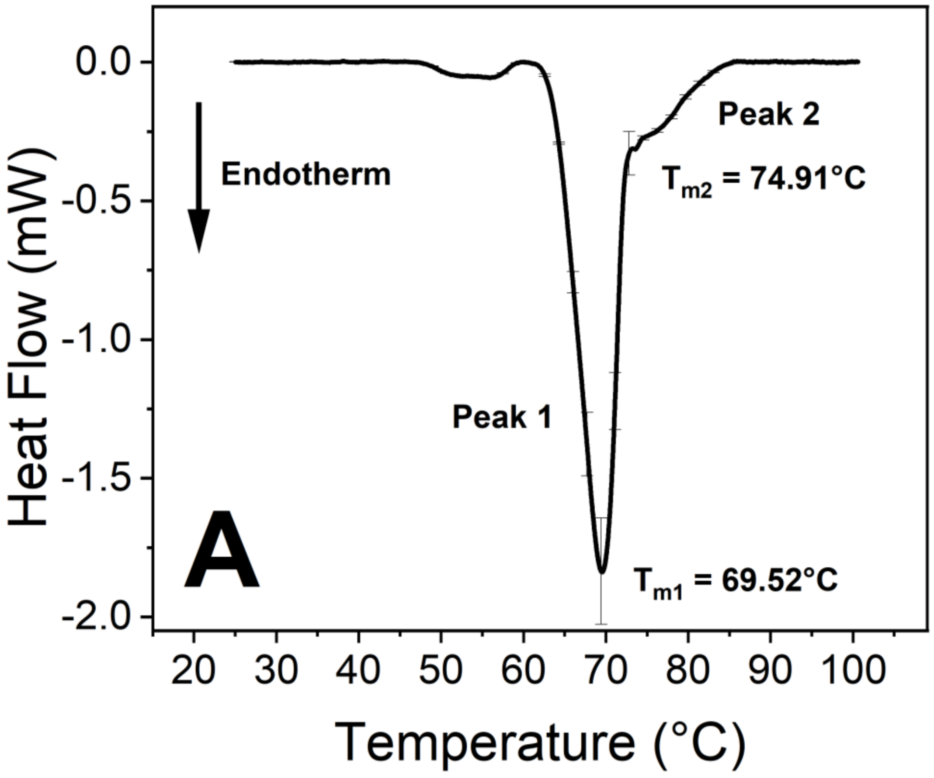


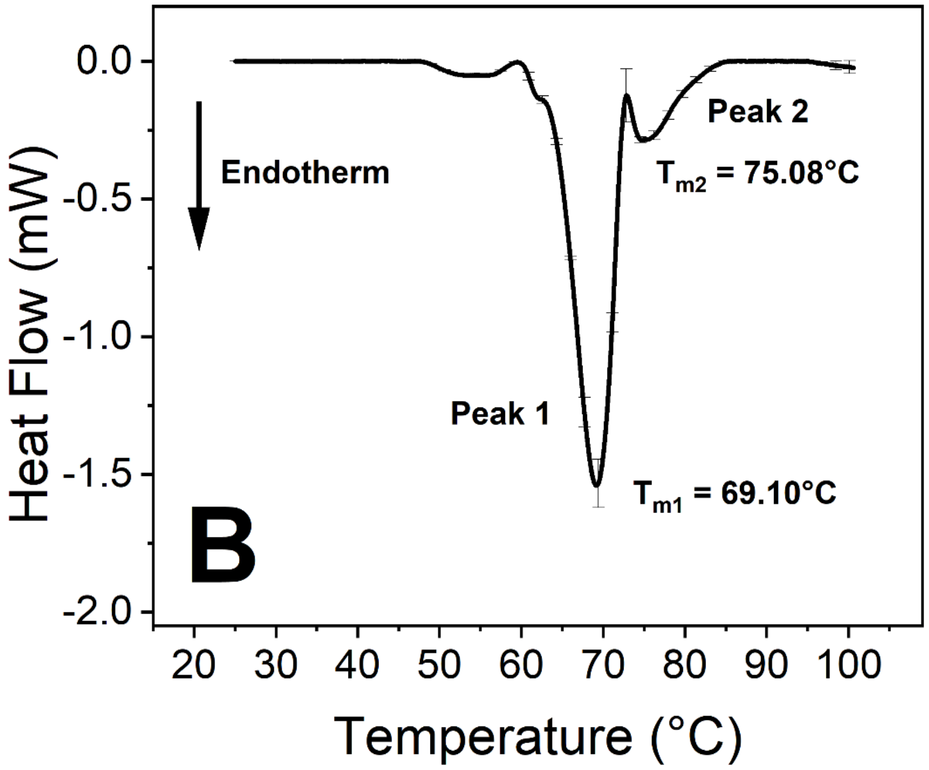


**Figure S3:** Thermal analysis of control (A) and virus-treated (B) anticoagulated whole blood samples after 15 hours of incubation at 37 °C. The DSC data represent the mean ± SD of four independent measurements (n = 4).
